# Probing weak lipid-lipid binding domain interactions with a mechanically transduced immunosorbent assay (METRIS)

**DOI:** 10.1101/2022.08.22.504198

**Authors:** Joshua P. Steimel, Joe Sarkis, Benjamin R. Capraro, Tom Kirchhausen, Alfredo Alexander-Katz

**Affiliations:** Department of Mechanical Engineering, University of the Pacific, Stockton, CA 95211, USA; Department of Cell Biology, Harvard Medical School, Boston, MA, 02115 USA; Program in Cellular and Molecular Medicine at Boston Children’s Hospital, Boston, MA 02115, USA; Department of Materials Science and Engineering, Massachusetts Institute of Technology, Cambridge, MA, 02139, USA

**Keywords:** METRIS, Rolling parameter, Ferromagnetic, Friction, Lipid Bilayer, K_*d*_, PTEN, Auxilin 1, GAK, endocytosis

## Abstract

Protein-lipid interactions constitute a very important class of biological interactions critical for multiple cell and tissue functions. It is believed that most lipid-protein interactions are very weak, with affinities in the 1mM-1*μ*M range. Here we study the interactions of multiple protein lipid binding domains with lipid membranes containing signaling lipids known as phosphatidylinositol phosphates (PIPs) using a new mechanically transduced immunosorbent assay (METRIS). We demonstrate that this assay can measure extremely weak interactions at PIP bilayer concentrations below 1%, which is close to the biological lipid concentration regime. In particular, we have studied the interaction of DrrA_*WT*_, DrrA_*K*568*A*_, PH-*δ*, and 2XFYVE as well as previously unexplored lipid binding domains such as Auxilin 1 (PTEN) and Auxilin 2 (GAK) against a wide palette of PIPs. Our results confirm that each of these domains interacts specifically with a PIP partner. In the case of Auxilin 1 and Auxilin 2, both proteins in the Clathrin endocytotic pathway, we find that their PTEN-like domain interacts specifically yet with ultra low affinity with PI3P and PI4P respectively. We have also found a new unknown medium-high affinity interaction between GAK with PI34P2. Our work, thus, provides a direct and robust method to measure and catalog protein lipid interactions which are important in many processes such as signaling and membrane sculpting. Furthermore, this assay can be extended in a straightforward manner to study other interactions such as ligand-receptor or antibody-antigen.

## Introduction

Lipid-protein interactions play an important role in multiple processes inside and outside the cell. In many circumstances, the interactions between lipids and proteins are very weak with affinities above 1*μ*M, yet they tend to be rather specific for a particular lipid-protein pair. Many of these lipid-protein interactions are related to lipid signaling and membrane sculpting [9,16,17,20]. One such example of lipid-protein interaction are kinases and phosphatases which can phosphorylate or dephosphorylate phosphatidylinositol lipids (PIP) that serve as signaling agents in multiple cellular processes and pathways [4,5]. It is believed that such interactions need to be weak yet specific because these protein enzymes must catalyze their reactions in multiple substrates within a short amount of time [11]. Otherwise their catalytic function would be very inefficient. In other cases, like endocytosis, the overall process is short lived, on the order of 100s of milliseconds to several seconds, which imposes a bound on the lifetime of the protein-lipid interactions occurring during endocytosis. Furthermore, the complete process includes multiple steps where proteins on the lipid membranes must exchange, implying that their lifetime must be shorter that the time it takes to complete this process. These are only some examples, yet it is believed that weak interactions between proteins or between lipids and proteins are at the core of a myriad of processes that drive the most dynamic aspects of cellular and tissue function [6,7,15].

The weak affinity of important lipid-protein interactions implies a short lifetime of the bonds as well as equilibrium constants in the order of tens of *μ*Ms, which makes studying such interactions with traditional methods such as ELISA impossible. Other methods such as Surface Plasmon Resonance and Layer Interferometry have been used with mixed success in weak lipid-protein interactions due to the difficulties associate with managing large quantities of the reagents, preparation of the lipid surfaces, and the need to have high density of binding sites in such substrates.

To overcome such limitations, here we study the interactions of several lipid binding domains with PIP lipids using a new mechanically transduced immunosorbent assay (METRIS) [1,2,21,24]. METRIS leverages the fundamental physical concept of friction and utilizes rolling protein-functionalized ferromagnetic beads of diameter D on a supported lipid bilayer (SLB), as seen in Fig. 1A-C.

**Figure 1.**
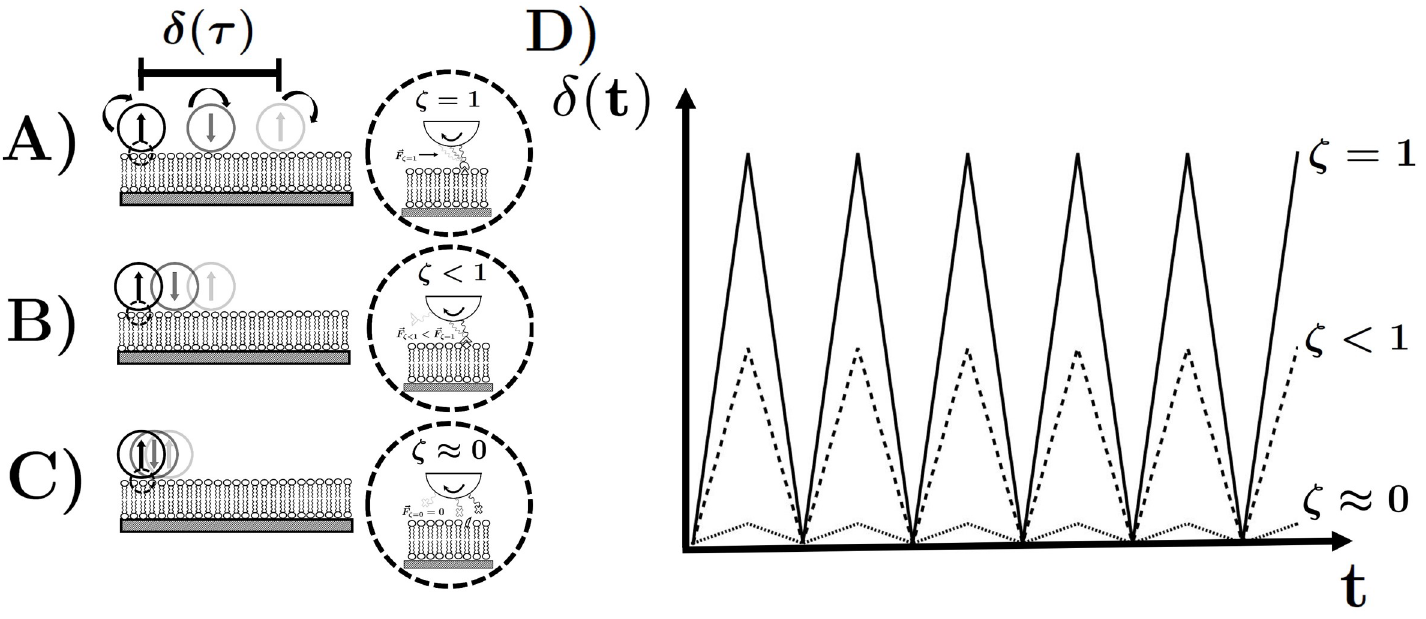
Schematic of mechanically transduced immunosorbent assay. A) Scenario of perfect rolling. The roller functionalized with the biological lipid binding domain *grabs* the substrate coated with the corresponding ligand associates with the ligand until eventually the bond breaks and forms another bonding with the leading edge of ligands. This interaction increases the effective friction between the substrate and the roller leading to, in this scenario, perfect conversion of rotational to translational motion. B) Intermediate rolling scenario. The lipid binding domain on the roller interacts with the ligand on the substrate but the affinity is lower than the scenario of perfect rolling. The effective friction is reduced and less rotational motion is converted into translational motion. C) Scenario of perfect slipping. There is no interaction or association between the lipid binding domain and the ligand on the substrate so the effective friction is zero in this case. The roller does not convert any of the rotational motion into translation. However, with this technique there will always be the contribution of hydrodynamic friction to the effective friction, so this scenario is not observable.

Using this new assay we are able to study such weak interactions under conditions similar to those occurring in biological settings, for example the concentration of PIPs in a PC membrane that could be below 1%. Also, METRIS only uses extremely small quantities of reagents, in particular proteins or lipid binding domains, something that might be useful when it is difficult or expensive to obtain large quantities of the protein in question. The method consists of rolling beads at a constant frequency, *ν*, by applying a torque using a rotating magnetic field. The frequency, *ν*, corresponds to the rotational frequency of the magnetic field. The field is applied using a series of coils as seen Fig. 8 The read-out is the rolling parameter *ζ*

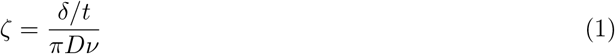

which is the ratio of the measured translational velocity, *δ/t*, to the velocity of a perfectly rolling bead without slip given by *πDν. δ* corresponds to the displacement of the bead in time, t, as seen in Fig. 1D for different rolling parameter values. If the rolling parameter is 1, the bead does not slip and implies a strong interaction between the lipid binding domain and the substrate. As the bead rolls, the bond between the proteins and the lipids on the substrate lead to an effective friction force on which the bead *grabs* the substrate, as depicted schematically in Fig. 1A. Eventually the protein-lipid pair must break, but newer ones are formed from the front of the contact zone. If the lipid-protein interactions are weaker, the bonds do not last as long and thus the effective friction force on the beads is reduced leading to a rolling parameter below 1 as seen in Fig. 1B. The limiting case of no interactions yields experimentally a finite but very small rolling parameter as seen in Fig. 1C. The origin of the friction in this case is hydrodynamic and is always present. This corresponds to the base line. Several experimental videos showing the different translation induced by binding events can be seen in Fig. 9. The basic premise of METRIS is that friction between a rotating bead and a surface is dependent on the interaction strength between the protein with which the beads are functionalized and the substrate.

## Materials and Methods

All lipids were supplied by Avanti Polar Lipids: 18:1 PC (cat. No. 850375), Liver L-α-phosphatidylinositol (cat. No.840042), 18:1 PI(3)P (cat. No. 850150), Brain PI(4)P (cat. No. 840045), 18:1 PI(5)P (cat. No.850152), 18:1 PI(3,4)P2 (cat. No. 850153), 18:1 PI(3,5)P2 (cat. No. 850154), Brain PI(4,5)P2 (cat. No. 84ü04β), 18:1 PI(3,4,5)P3 (cat. No. 850156), 18:1 Biotinyl Cap PE (cat. No. 870273).

### 0.1 DNA Constructs

Residues 40-400 of auxilin1 (B. taurus; hereafter referred to as PTEN-Aux1) were inserted into a pRSET-based vector also encoding a N-terminal His6-tag followed by a TEV cleavage sequence. A DNA construct encoding residues 400-766 of GAK (H. sapiens; hereafter referred to as PTEN-GAK) as an N-terminal His6-tagged, SUMO (small ubiquitin-like modifier)-fusion was generated by PCR followed by ligation-independent cloning into a non-commercial vector based on the pET series that features a T7 promoter for inducible bacterial expression. The PH domain of PLC*δ* was inserted into a pGEX-4T-based vector modified to include a TEV cleavage site. DNA encoding residues 340-647 of DrrA (L. pneumophila; wild-type and K568A point mutation) was inserted into pET-19. All DNA constructs were verified by DNA sequencing. A plasmid encoding the FYVE domain of Hrs as a His6-tagged GST fusion protein was obtained from M. Lemmon, Yale University School of Medicine.

### 0.2 Protein Preparation

The PH domain of PLC*δ* was purified and cleaved from GST following published procedures. PTEN-Aux1, WT and K568A DrrA, GST-FYVE, and PTEN-GAK, were expressed in BL21 E coli. Cells were lysed using sonication. Lysates were clarified by centrifugation, and the supernatant was applied to nickel-NTA resin. Proteins were eluted with imidazole. His6-tags and any fusion protein moieties were not removed for lipid-binding studies, with the exception of PTEN-Aux1 and the PH domain of PLC*δ*, which were treated with TEV protease for tag removal. Proteins were dialyzed into bicarbonate buffer and biotinylated using EZ-Link^*TM*^ Sulfo-NHS-LC-Biotin (ThermoFisher) following the manufacturer’s protocol. For some preparations, an HABA/avidin assay (ThermoFisher) was used to quantify the extent of biotinylation (3-4 biotin/protein).

### 0.3 Supported Lipid Bilayer Preparation

Lipid Mixture, formed by 90*mol* : *mol* of DOPC and 10*mol* : *mol* of specific phosphoinositide, was solubilized in chloroform/methanol/water (20 : 9 : 1v/v/v) and thoroughly mixed in a glass tube for a total amount of 0.3mg. Lipids were adhered along the sides of the glass tube under a stream of nitrogen gas by gently rotating the tube until the solvent had evaporated. Residual traces of solvent were then evaporated under vacuum overnight. 300 *μ*L of Lipid Buffer (Hepes 20mM, NaCl 150mM pH 4.8 or pH 6.8; acidic pH was used to improve the homogeneity of the distribution of PIs in the bilayer 5) was added to the dried lipid, and the tube was allowed to incubate at room temperature for 45-60 mins. This incubation allows the lipids to gradually peel off the glass surface and swell. Multiple vigorous vortexing cycles were applied, forming vesicles with heterogeneous size. Small unilamellar vesicles at 1mg/mL were formed by 31 extrusion cycles using polycarbonate membranes of 50nm pore size (Avanti Polar Lipids). This solution was used directly to form the supported lipid bilayer.

Microfluidic channels were created as the support for the lipid bilayer. Two pieces of doubled sided tape, provided by 3M, were placed on glass slides, provided by Fisher Scientific. A cover slip, provided by VWR, was placed on top of the double sided tape creating a channel approximately 22mm × 5mm. Once the channel was created a solution of NaOH was pipetted into the channel and left to sit for approximately 5 minutes. The solution was then flushed with a lipid buffer solution, pH 4.8, by pipetting the solution in from one side of the channel and sucking it out from the other end with a kimwipe. The channel was washed several times to remove a NaOH residue. 25*μ*l of the lipid stock solution, at 1mg/mL, was diluted in 250μL of the 4.8 pH buffer solution. The diluted solution of lipids was then inserted into the channel allowed to sit for 30 minutes. The channel was monitored throughout to ensure no evaporation took place which can disrupt the lipid bilayer formation. After this 30 minute period the channel was washed again several times with a 6.8 pH buffer solution, making sure no evaporation occurred in the channel. 10*μ*L of the desired functionalized bead solution, described in the section above, was diluted into 800*μ*L of the 6.8 pH lipid buffer. This diluted solution of beads was then inserted into the channel and the channel was sealed at both ends with epoxy to prevent evaporation. The channel was then placed on top of neodymium magnet to magnetize the ferromagnetic particles and then placed on the slide holder within the experimental apparatus.

### 0.4 Ferromagnetic Roller Functionalization

Streptavidin coated ferromagnetic particles, 9*μ*m in diameter and provided by Spherotech, were diluted in an aqueous solution from a 4mL 1.0% w/v stock. 100*μ*L of the stock solution was diluted into 5mL of water. 1mL was extracted from the diluted solution and the desired amount of biotinylated protein was added to the solution to coat streptavidin coated ferromagnetic particles. To ensure that all the sites were completely coated with protein the amount of protein added was enough to coat the beads 50×. The bead and protein solution was then left to react at 4°C overnight.

## Results and Discussion

### 0.5 METRIS Measurement of Biotin-Streptavidin Interaction on Supported Lipid Bilayer

By comparing between different rolling parameters from the same beads on different substrates we are able to discern very weak yet specific interactions. To test this technique, we first studied the canonical interaction between biotin and streptavidin. The streptavidin coated beads were rolled on a biotin functionalized glass slide, provided by Xenopore, which was then coated with avidin ligands. The displacement was measured as a function of time for different ratios of avidin to biotin binding sites on the substrate, as illustrated in the kymographs in Fig. 2A. The measured rolling parameter was approximately 0.95 for an unpassivated biotinylated substrate as seen in Fig. 2B. This value intuitively makes sense as the biotin-streptavidin is the strongest non-covalent interaction, K_*d*_ ≈ 10^−15^M. The beads are sensitive enough to measure the full biotin-streptavidin interaction until all of the available biotin binding sites are coated with avidin binding sites and the hydrodynamic friction limit is not reached until the avidin to biotin ratio reaches 20. The biotin-streptavidin interaction was then measured on a PIP lipid bilayer with 10% of that bilayer being composed of biotinylated lipids. As seen in Fig. 2B, the strength of the biotin-interaction on the lipid bilayer substrate is lower than measured for the glass substrate however this is most likely due to the mobility of the biotinylated lipids and the fact that the beads might be stripping lipids from the membrane rather than breaking from the biotinylated lipids. This can also explain why at longer times the interaction seems to be reduced.

**Figure 2.**
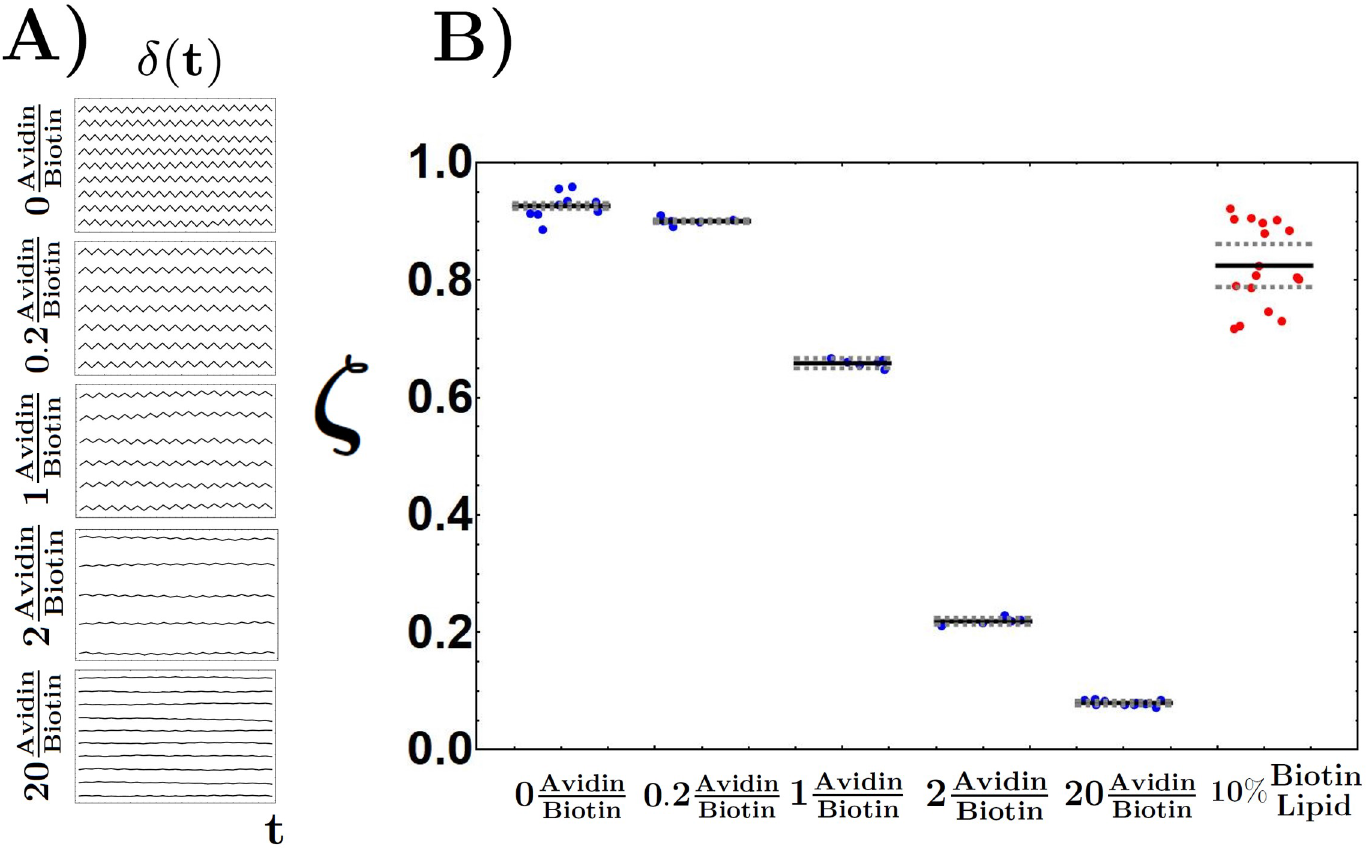
Rolling parameter for biotin-streptavidin interaction on avidin substrate and a supported lipid bilayer. A) Kymographs of rollers functionalized streptavidin rolling on a biotin coated substrate passivated with increasing amounts of avidin. B) The rolling parameter for an unpassivated surface, no avidin molecules, is approximately 0.95. However, as the ratio of avidin to biotin binding sites increase the rolling paramter decreases as there are less available binding sites. The supported lipid bilayer contains 10% biotinylated lipids. The rolling parameter appears to be lower for the supported lipid bilayer than the biotin glass substrate due to biotin lipids being pulled from the lipid bilayer. The solid black line indicates the average rolling parameter of the entire data set. The dashed grey lines represent the 95% confidence interval of each dataset.

### 0.6 METRIS Measurement of DrrA_*WT*_, DrrA_*K*568*A*_, PH-*δ*, and 2XFYVE with PIPs

Other well known proteins that interact with different PIP lipids with a varying degree of affinity were also characterized. In particular, we studied DrrA_*WT*_, DrrA_*K*568*A*_, PH-*δ*, and 2XFYVE interacting with PC, PIP, PI3P, PI4P, PI34P_2_, and PI45P_2_. The lipids chosen were selected on the basis of the presence and position of the phosphate group on the inositol ring as well as the degree of overall charge. As can be seen from Fig. 3, our results confirm previous studies [3,8,12,18,22,23], which had shown that these set of binding domains having specific interactions with a particular PIP. Moreover, the relative affinities of the interactions are in agreement with the obtained rolling parameters, higher affinity interactions exhibit larger rolling parameters.

**Figure 3.**
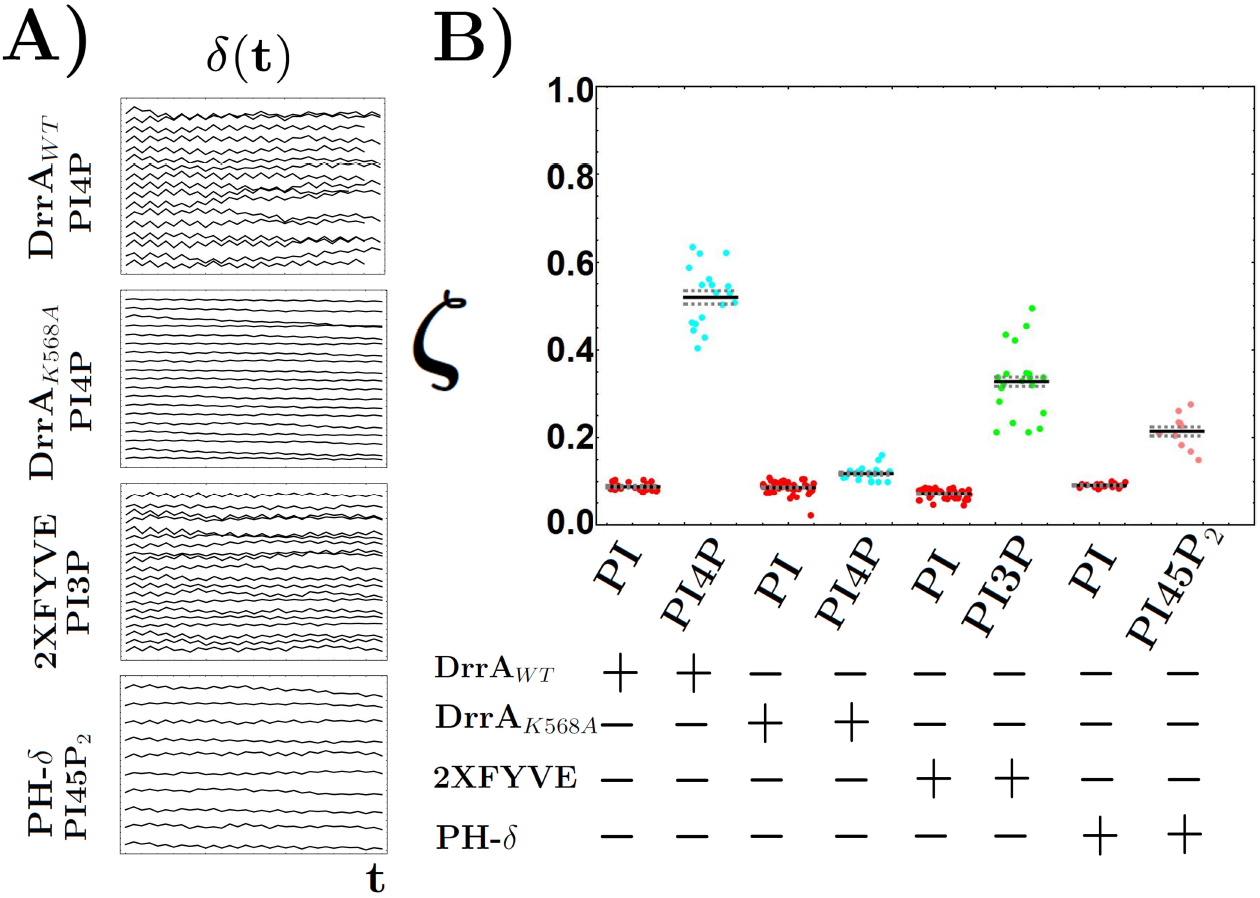
Rolling parameter for interactions on supported bilayer with known binding affinity. A) Kymographs of rollers functionalized with DrrA_*WT*_, DrrA_*K*568*A*_, PH-*δ*, and 2XFYVE rolling on supported lipid bilayers (SLB) containing 10% PI4P, PI4P, PI3P, and PI45P_2_, respectively. The oscillations in motion are large for DrrA_*WT*_ which is to be expected due its high affinity for PI4P but the oscillations become much smaller and the translation diminishes drastically once a point mutation is introduced, DrrA_*K*568*A*_. Similarly the amplitude of oscillations decrease from 2XFYVE to PH-*δ* as expected. B) The rolling parameter is plotted for different biological interactions with SLBs. Each point is the average rolling parameter measured for a roller, there are 34 measurements per roller. The solid black line indicates the average rolling parameter of the entire data set. The dashed grey lines represent the 95% confidence interval of each dataset. The SLB contains 10% of each ligand. The rolling parameter correlates well with the known binding affinity of these interactions and is specific to the ligand phosphorylation.

A graphical image showing the displacement of at least ten different beads rolling back and of forth on multiple substrates is shown in Fig. 3A. The frequency of rolling is set to 1 Hz unless otherwise noted. In the top kymograph we show the rolling of DrrA_*WT*_ coated beads on membranes containing 10% of PI4P. As can be seen from the amplitude of oscillation, the rolling parameter in this case is high, which is expected as DrrA_*WT*_ exhibits a high affinity for PI4P, 5nM. By introducing a point mutation into the DrrA_*WT*_ protein we can see this strong affinity for PI4P is lost in the mutated DrrA_*K*568*A*_ protein as the oscillations are now much smaller. The same can be seen for 2XFYVE and PH-*δ* on SLB containing 10% PI3P and PI45P_2_ respectively. The oscillations are smaller for PH-*δ* which is expected as the affinity for PI45P_2_ is lower than the affinity of 2XFYVE for PI3P [3,12,18,22]. Interestingly it is apparent that the oligomeric state of the protein appears to have no effect on the rolling parameter measurement as seen here and moving forward.

A convenient way to quantify the relative strength of the interaction just mentioned before is to plot the average rolling parameter and 95% confidence interval for each system. This is shown in Fig. 3B. The interaction affinity of DrrA_*WT*_ for PI4P, 2XFYVE for PI3P, and PH-*δ* for PI45P_2_, have been previously reported and to be approximately 5nM, 100nM, and 1*μ*M respectively [3,8,12,18,22,23]. As we see in Fig. 3B the rolling parameter seems to match well qualitatively with those literature values and confirms that this new technique can resolve these previously investigated interactions. These measured rolling parameters are also much larger than when rolled on the PI SLB indicating the specificity of these interactions.

### 0.7 METRIS Measurements of DrrA_*WT*_, DrrA_*K*568*A*_, PH-*δ*, and 2XFYVE with PIPs are Stereospecific

We further illustrate that this rolling bead assay can clearly identify the stereospecificity of these known interaction as seen in Fig. 4. Here we see that DrrA_*WT*_ is specific to PI4P, 2XFYVE to PI3P, and PH-*_δ_* to PI45P2 compared to other functionalized SLBs. We also explored the interaction between PIPs and two proteins Auxilin 1 and Auxilin 2 that are required in the endocytotic pathway. Auxilin 1 has a PTEN-like domain and Auxilin 2 is another protein that is more prevalent in the brain and believed to be very important for brain functions in synaptic joints [10, 19]. The PTEN of Auxilin has been showed to exhibit an affinity for certain PIPs however determining the relative affinities between auxilins has been difficult using traditional assay techniques like lipid dot assay, spin down assays, or ELISAs. Here we compare the relative affinities of Auxilin 1 and Auxilin 2 to specific PIPs, with Auxilin 1 exhibiting a preference for PI3P and Auxilin 2 exhibiting a preference for PI4P. Previous studies have investigated Auxilin 1 and 2 affinities for PIPs independently. Lipid-dot assays performed on the GST-PTEN Auxilin 2 or GAK, the dimer, illustrated an affinity preference of *PI*3*P*=*PI*4*P*=*PI*5*P* > *PI*35*P*2 >*PI*45*P*_2_. Due to the intrinic dimer of Auxilin 2 this assay has difficulty in differentiating between the affinity of PI3P and PI4P. [10] In the case of the full length version of Auxilin 1 lipid dot assays with full length monomer Auxilin 1 exhibited an affinity ranking of *PI*3*P* > *PI*4*P* = *PI*34*P*_2_> *PI*45*P* = *PI*45*P*_2_. [19] Additionally, spin down lipid vesicle assay measurements have shown that the monomer PTEN-Auxilin1 exhibited an affinity of *PI*4*P* > *PI*45*P*_2_. [13]. The issue with these experiments is these experiments require high concentration of PIPs in the vesicle, approximately 15%, higher than the 10% used in this study, due to the tendency of protein washing away during centrifugation. This issue is only exacerbated by the low affinity of the interactions being measured. Total internal reflection florescence (TIRF) have also indicated this binding preference [14]. Here we have measured the relative affinity of PTEN monomer Auxilin 1 and Auxilin 2 GAK for several PIPs including PC, PI, PI3P, PI4P, PI5P, PI35P_2_, PI34P_2_, PI45P_2_, and PI345P_3_in a robust, reproducible, simple, and high throughput methodology using METRIS which requires minimal material and can also be used to estimate the binding affinity of the interaction.

**Figure 4.**
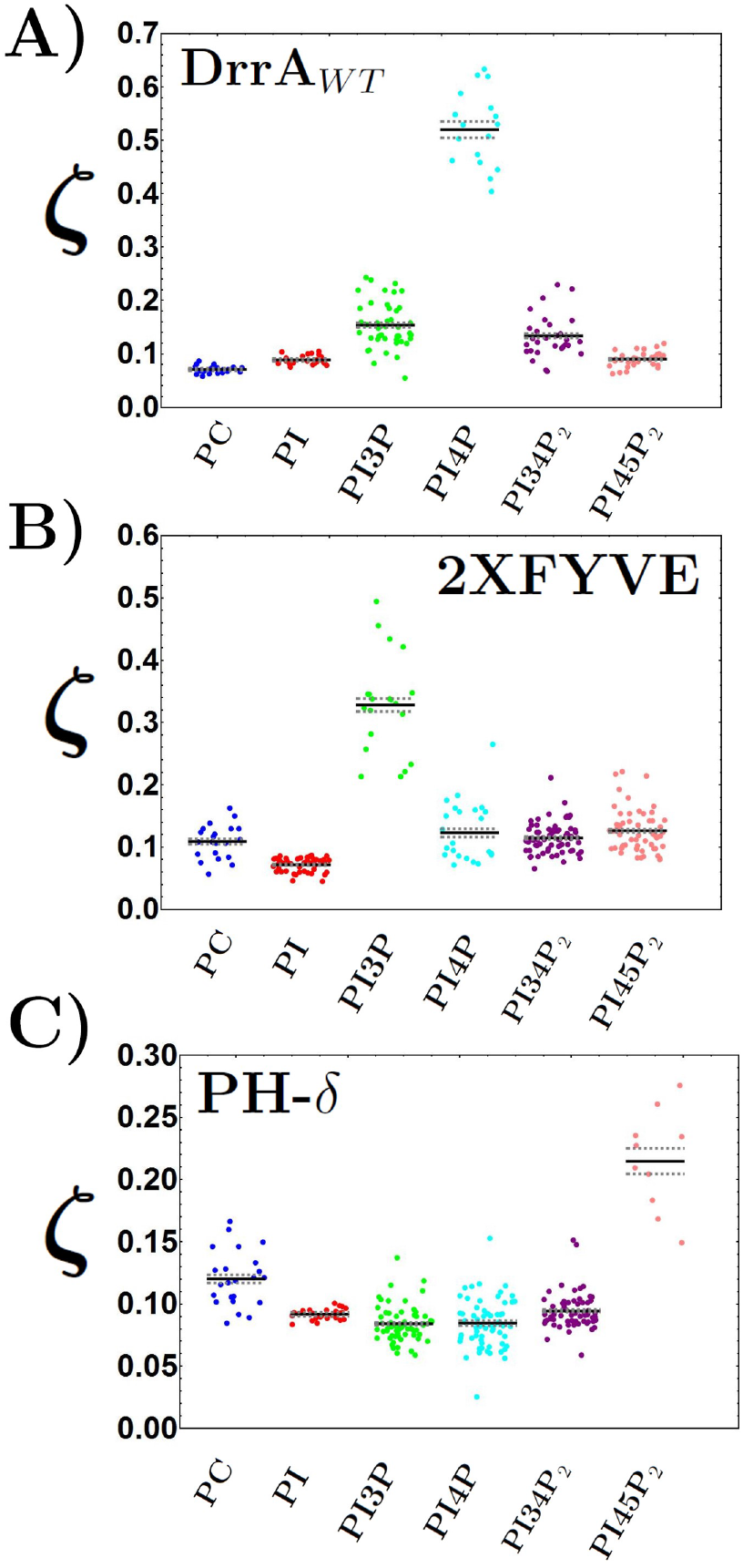
Roller binding assay illustrates stereospecificity of biological interactions with SLB. A) Rolling parameter of DrrA_*WT*_ on SLB with 10% PC, PI, PI3P, PI4P, PI34P_2_, and PI45P_2_. DrrA_*WT*_ shows extremely high affinity for PI4P compared to other SLB functionalization’s. B) Rolling parameter for 2XFYVE shows preferential affinity for PI3P. C) Rolling parameter for PH-δ show preferential affinity for PI45P_2_.

### 0.8 METRIS Confirms PTEN Stereospecific Preference for PI3P and GAK for PI4P

As can be seen from the rolling parameters seen in Fig. 5, we confirm that the PTEN-like domain of Auxilin 1 and Auxilin 2 are specific for PI3P and PI4P respectively which is consistent with the literature data. Moreover by comparing the rolling parameter values it is clear that the PTEN Auxilin 1 prefers PI3P over PI4P and PI45P_2_ whereas Auxilin 2 or GAK prefers PI4P, based on the relative affinities, and an unexplored interaction even larger than PI4P, PI34P_2_ over both PI3P and PI45P_2_. Here we speculate that GAK-PI34P_2_ interaction is stabilized cooperatively by charge. Thus, this technique is able to distinguish between relativity affinities between PIP interactions using less protein material and PIP concentration than other traditional techniques and is sensitive enough to measure and differentiate between these weaker interactions.

**Figure 5.**
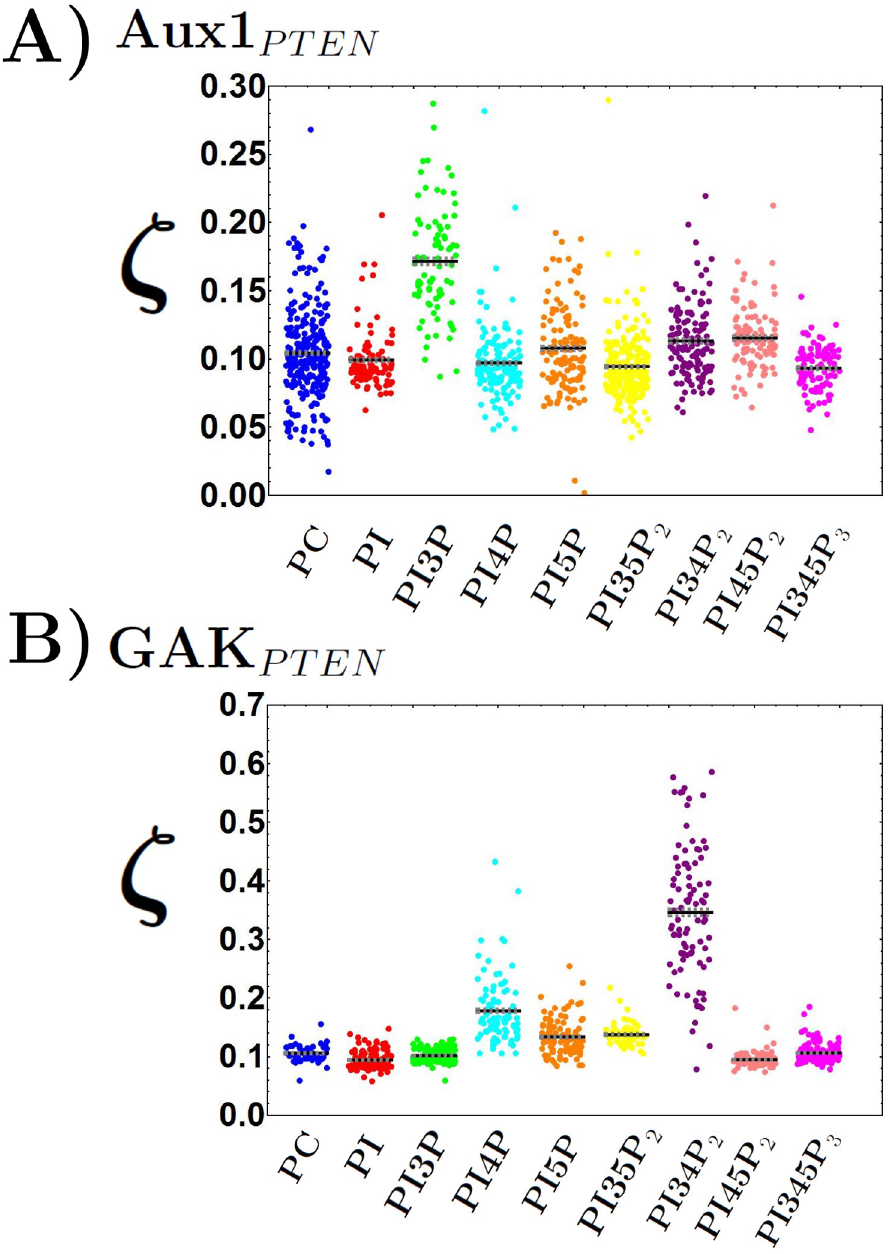
Roller binding assay shows preferential affinity of Aux1_*PTEN*_ for PI3P and GAK_*PTEN*_ for PI34P_2_ and PI4P. A) Rolling parameter measurements show Aux1_*PTEN*_ affinity for PI3P when compared to other PIPs. This correlates well with biological observations. B) Rolling parameter measurements show GAK_*PTEN*_ affinity for PI4P and a newly discovered interaction with PI34P_2_. Again this result correlates well to biological observations.

### 0.9 METRIS Assay Ultrasensitive and Measures Binding at Biological Concentrations

To further demonstrate that this technique is ultrasensitive, we present in Fig. 6A a plot of the rolling parameter as a function of the degree of functionalization of the bead, as well as the PIP content in the lipid bilayer for 2 different proteins DrrAW T and GAK as seen in Fig. 6B-C. As can be seen from the plot, the rolling parameter exhibits a marked drop when the density of protein in the surface is below the theoretical maximum loading, yet the signal is saturated once the amount of protein used for functionalization is above the theoretical limit. Perhaps more importantly, the sensitivity remains constant as one decreases the concentration of PIP lipid in the membrane from 15% to 1%. These results indicate that this technique might be equally sensitive all the way to concentrations of 0.1%, which puts us squarely in the biological concentration realm.

**Figure 6.**
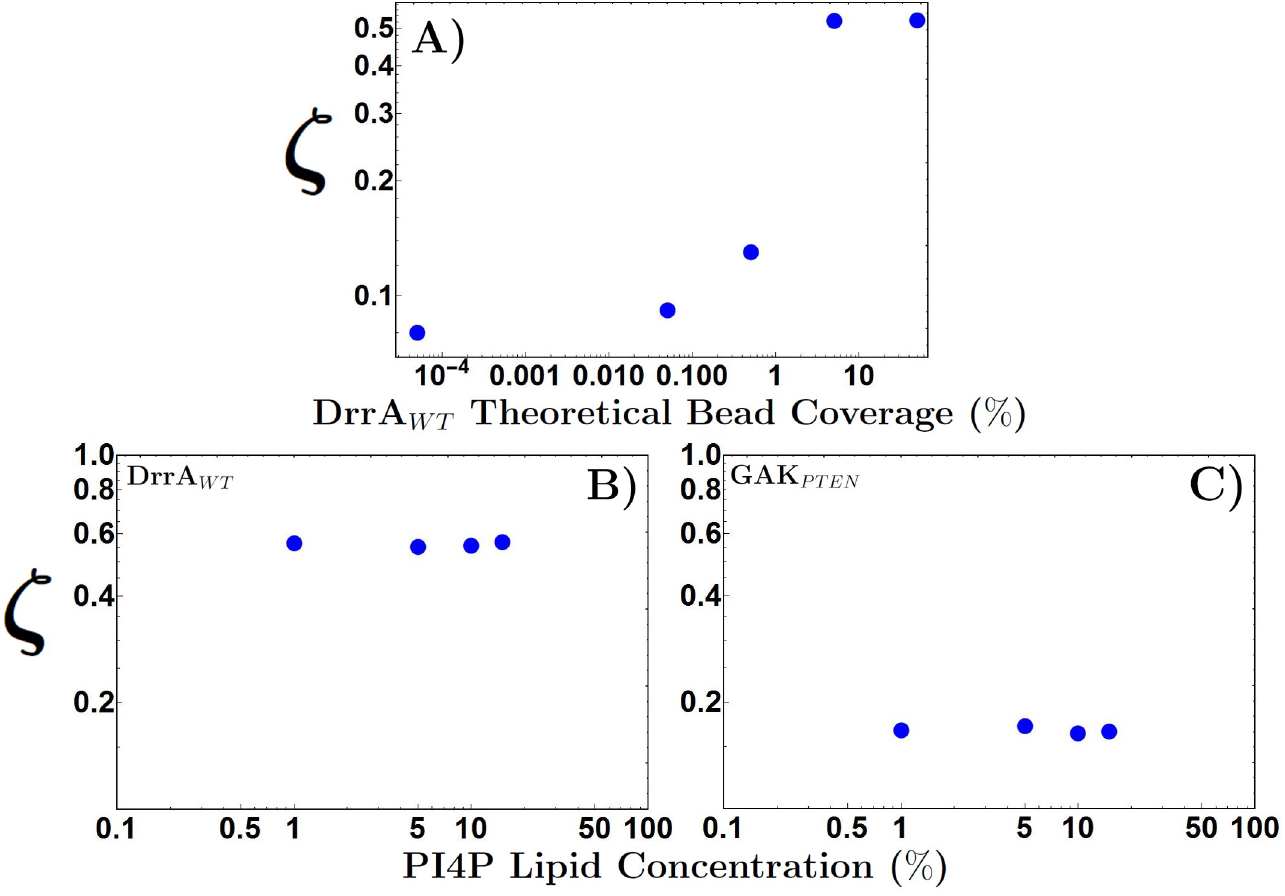
Sensitivity of rollers to concentration of protein and lipids. A) Rolling parameter decreases as concentration of DrrA_*WT*_ on the surface of the rolling bead is below the theoretical maximum loading capacity. The lipid bilayer is composed of 10% of PI4P lipids. B) Rolling parameter as a function of the concentration of PI4P lipids for a roller coated with DrrA_*WT*_ and C) GAK_*PTEN*_. The rollers are clearly able to discern differences in effective friction up to 1% PI4P concentration.

### 0.10 METRIS Provides Estimate of Binding Affinity of Lipid Binding Domains and PIPs

As demonstrated, METRIS provides a robust method to screen weak interaction to determine relative affinities between interactions. However, METRIS can provide quantitative measurements of binding affinity, *K_d_*. Rolling parameter values were measured for DrrA_*WT*_-PI4P, 2XFYVE-PI3P, and PH-*δ*-PI45P_2_ which have been well studied and known *K_d_* values. One can then fit a Log-Log plot of the measured rolling parameter for known *K_d_* values, including the biotin-streptavidin interaction which has a binding affinity of approximately 10^-15^M and a null interaction of DrrA_*WT*_-PI which was given an estimated *K_d_* of approximately 1M. This fitted function allows one to estimate binding affinity values for previously unknown interactions as previously demonstrated [1,2,21,24]. With other traditional optical readout based techniques determining the preference in binding affinity of Auxilin 1 to PI4P and Auxilin 2 to PI3P was either not possible or extremely difficult. With METRIS the rolling parameter readout demonstrated a clear binding affinity preference but when extrapolating the value the measured *K_d_* shows that Auxilin 1 and PI3P have a binding affinity of approximately 144*μ*M and Auxilin 2 and PI4P exhibits a binding affinity of approximately 84*μ*M. This is in stark contrast to the other PIP binding pair of Auxilin 1 and PI4P which exhibited a binding affinity of 0.3M and Auxilin 2 and PI3P was found to be approximatley 0.18M. This finding also demonstrates that this PIP interaction is indeed quite low and at the limit of other techniques but the METRIS technique was able to measure this difference in a clear and robust manner.

## 1 Conclusion

In summary, we have introduced a new robust technique based on rolling beads to study weak interactions. We have applied this assay to study previously known lipid binding domain-lipid interactions to validate its sensitivity. We have then studied unexplored lipid binding domain-lipid interactions between Auxilin 1 (PTEN) and Auxilin 2 (GAK) and PIP lipids where we have that these proteins display specific yet ultra-weak interactions with PIP lipids, and discovered a new medium/high affinity interaction between GAK and PI34P2. Our results also show that it is possible in a straightforward manner to use the rolling bead method to study in a scalable and high throughput fashion other weak interactions. This new technique presents clear advantages over traditional equilibrium assay techniques which utilize labeled ligands as an optical signal to indicate binding events. The amount of proteins or ligands utilized with this method is very small, picomolar, compared to other equilibrium methods which require concentration close to *K_d_* concentrations. This become problematic for any *K_d_* values above micromolar concentrations as proteins can tend to aggregate. Additionally, this technique is well suited for studying weak interactions where it is difficult or potentially impossible to produce proteins at micromolar concentration or more. Thus, we expect it to become an important tool in the discovery and characterization of weakly interacting complexes in biology.

**Figure 7.**
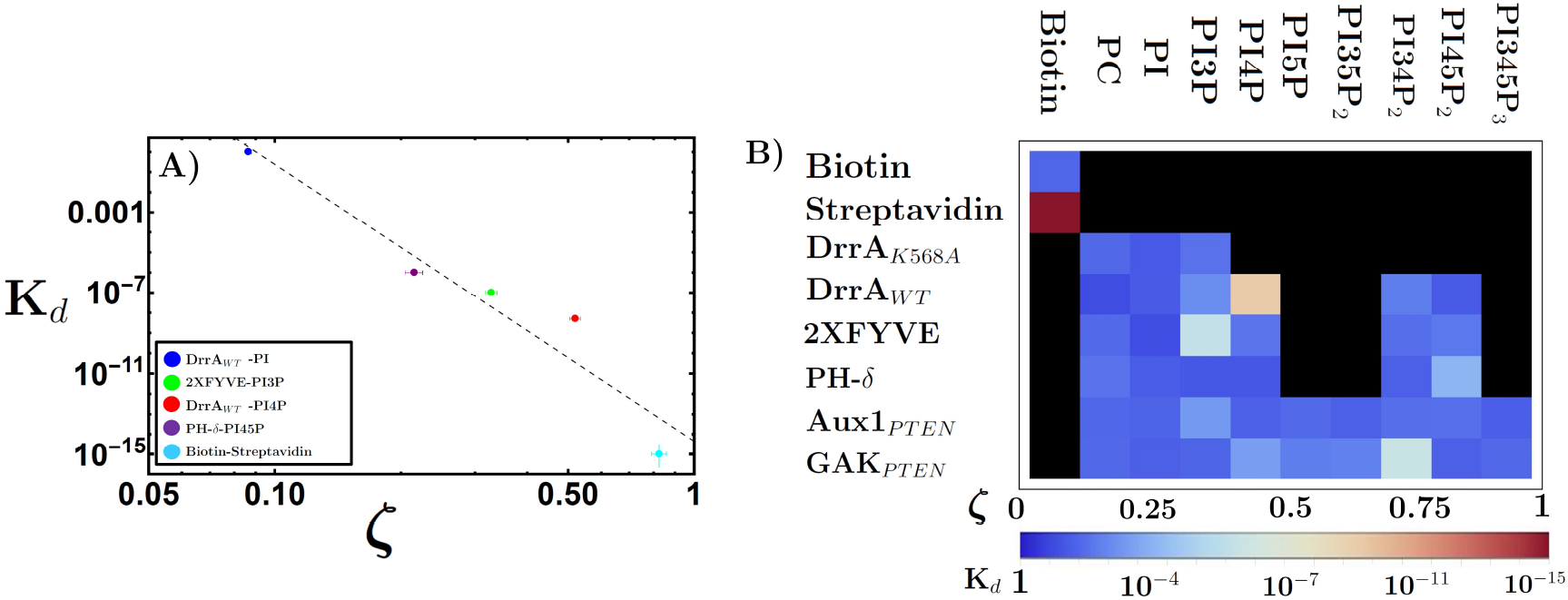
Estimation of relative binding affinity of lipid binding domain-lipid interactions. A) Log-Log plot of the rolling parameters (*ζ*) with reported *K_d_* values. The best fit line has an *R*^2^ correlation of approximately 0.9. Extrapolated binding affinity values of Auxilin 1 show a clear preference for PI3P, approximately 144*μ*M,compared to P14P, approximately 0.3M. Additionally, Auxilin 2 shows a clear affinity preference towards PI4P, approximately 84*μ*M, compared to PI3P, approximately 0.18M. These approximate *K_d_* values illustrate the strength of the METRIS technique to allow for the detection of weak interactions in ranges that are difficult for other optical readout techniques. This binding affinity preference correlates well observations in previous studies. B) Arrayplot of all of the binding affinities and associated rolling parameters for the interactions described.

## Supporting information

Example Video

## Acknowledgments

We thank M. Lemmon, Yale University, school of Medicine for sharing the FYVE domain plasmid. We thank Iris Rapoport for helping in protein preparation. We thank R. Marsland for facilitating this collaboration. This research was supported by NIH grant NIH R01 GM075252 (TK).

## Supporting Information

**S1 Figure**

**Figure 8.**
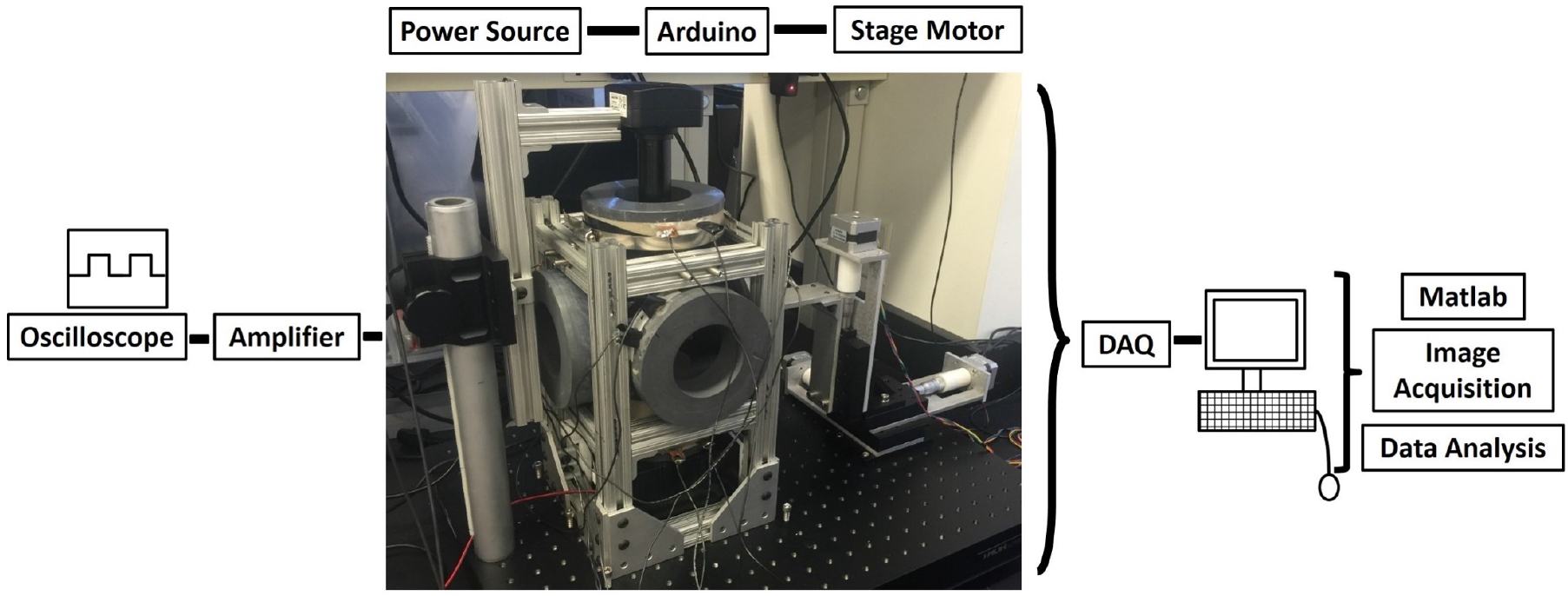
METRIS apparatus. Three pairs of Helmholtz coils were mounted on an aluminum T-slot assembly. Two sinusoidal signals are generated in Matlab, passed through a DAQ, amplifier, and then to the Helmholtz-like coils. Visualization is accomplished using a lens tube, 10X objective, and CCD camera. A 10 mT field was utilized.

**S2 Figure**

**Figure 9.**
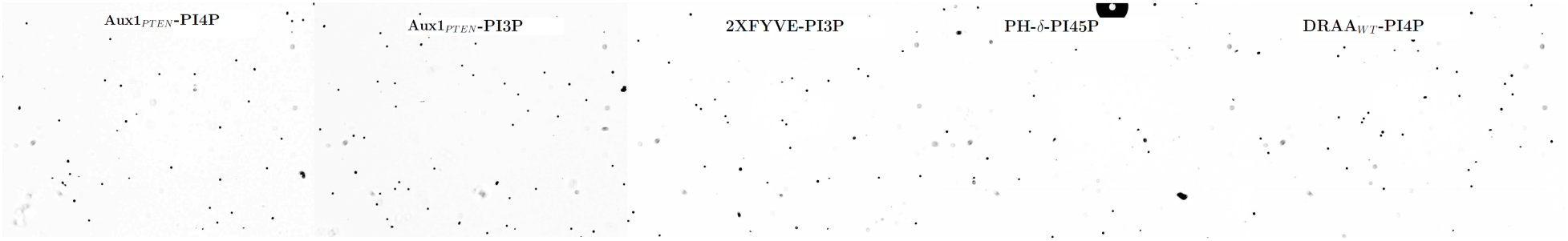
Videos of METRIS Measurements.

## References

1. US Patent for Systems and methods for detecting molecular interactions using magnetic beads Patent (Patent # 9,977,015 issued May 22, 2018) - Justia Patents Search.

2. Functional conservation and divergence of the helix-turn-helix motif of E2 ubiquitin-conjugating enzymes. The EMBO Journal, 41(3):e108823, Feb. 2022. Publisher: John Wiley & Sons, Ltd.

3. N. R. Blatner, R. V. Stahelin, K. Diraviyam, P. T. Hawkins, W. Hong, D. Murray, and W. Cho. The Molecular Basis of the Differential Subcellular Localization of FYVE Domains. Journal of Biological Chemistry, 279(51):53818–53827, Dec. 2004.

4. L. C. Cantley. The phosphoinositide 3-kinase pathway. Science (New York, N.Y.), 296(5573):1655–1657, May 2002.

5. L. C. Cantley and B. G. Neel. New insights into tumor suppression: PTEN suppresses tumor formation by restraining the phosphoinositide 3-kinase/AKT pathway. Proceedings of the National Academy of Sciences, 96(8):4240–4245, Apr. 1999.

6. P. Chien and L. M. Gierasch. Challenges and dreams: physics of weak interactions essential to life. Molecular Biology of the Cell, 25(22):3474–3477, Nov. 2014.

7. N. de Souza. The power of ‘weak’ interactions. Nature Methods, 13(1):14, Jan. 2016.

8. C. M. Del Campo, A. K. Mishra, Y.-H. Wang, C. R. Roy, P. A. Janmey, and D. G. Lambright. Structural basis for PI(4)P-specific membrane recruitment of the Legionella pneumophila effector DrrA/SidM. Structure (London, England: 1993), 22(3):397–408, Mar. 2014.

9. J. P. DiNitto, T. C. Cronin, and D. G. Lambright. Membrane recognition and targeting by lipid-binding domains. Science’s STKE: signal transduction knowledge environment, 2003(213):re16, Dec. 2003.

10. E. Eisenberg and L. E. Greene. Multiple roles of auxilin and hsc70 in clathrin-mediated endocytosis. Traffic (Copenhagen, Denmark), 8(6):640–646, June 2007.

11. B. H. Falkenburger, J. B. Jensen, and B. Hille. Kinetics of PIP2 metabolism and KCNQ2/3 channel regulation studied with a voltage-sensitive phosphatase in living cells. The Journal of General Physiology, 135(2):99–114, Feb. 2010.

12. F. M. Flesch, J. W. Yu, M. A. Lemmon, and K. N. J. Burger. Membrane activity of the phospholipase C-1 pleckstrin homology (PH) domain. Biochemical Journal, 389(2):435–441, July 2005.

13. R. Guan, H. Dai, D. Han, S. C. Harrison, and T. Kirchhausen. Structure of the PTEN-like region of auxilin, a detector of clathrin-coated vesicle budding. Structure (London, England: 1993), 18(9):1191–1198, Sept. 2010.

14. K. He, E. Song, S. Upadhyayula, S. Dang, R. Gaudin, W. Skillern, K. Bu, B. R. Capraro, I. Rapoport, I. Kusters, M. Ma, and T. Kirchhausen. Dynamics of Auxilin 1 and GAK in clathrin-mediated traffic. Journal of Cell Biology, 219(3):e201908142, Jan. 2020.

15. M. Y. Hein, N. C. Hubner, I. Poser, J. Cox, N. Nagaraj, Y. Toyoda, I. A. Gak, I. Weisswange, J. Mansfeld, F. Buchholz, A. A. Hyman, and M. Mann. A human interactome in three quantitative dimensions organized by stoichiometries and abundances. Cell, 163(3):712–723, Oct. 2015.

16. S. J. Leevers, B. Vanhaesebroeck, and M. D. Waterfield. Signalling through phosphoinositide 3-kinases: the lipids take centre stage. Current Opinion in Cell Biology, 11(2):219–225, Apr. 1999.

17. M. A. Lemmon. Phosphoinositide recognition domains. Traffic (Copenhagen, Denmark), 4(4):201–213, Apr. 2003.

18. M. A. Lemmon, K. M. Ferguson, R. O’Brien, P. B. Sigler, and J. Schlessinger. Specific and high-affinity binding of inositol phosphates to an isolated pleckstrin homology domain. Proceedings of the National Academy of Sciences of the United States of America, 92(23):10472–10476, Nov. 1995.

19. R. H. Massol, W. Boll, A. M. Griffin, and T. Kirchhausen. A burst of auxilin recruitment determines the onset of clathrin-coated vesicle uncoating. Proceedings of the National Academy of Sciences, 103(27):10265–10270, July 2006. ISBN: 9780603369100 Publisher: National Academy of Sciences Section: Biological Sciences.

20. Y. Nishizuka. Protein kinase C and lipid signaling for sustained cellular responses. FASEB journal: official publication of the Federation of American Societies for Experimental Biology, 9(7):484–496, Apr. 1995.

21. C. J. Petell, K. Randene, M. Pappas, D. Sandoval, B. D. Strahl, J. S. Harrison, and J. P. Steimel. Mechanically transduced immunosorbent assay to measure protein-protein interactions. eLife, 10:e67525, Sept. 2021. Publisher: eLife Sciences Publications, Ltd.

22. V. G. Sankaran, D. E. Klein, M. M. Sachdeva, and M. A. Lemmon. High-affinity binding of a FYVE domain to phosphatidylinositol 3-phosphate requires intact phospholipid but not FYVE domain oligomerization. Biochemistry, 40(29):8581–8587, July 2001.

23. S. Schoebel, W. Blankenfeldt, R. S. Goody, and A. Itzen. High-affinity binding of phosphatidylinositol 4-phosphate by Legionella pneumophila DrrA. EMBO reports, 11(8):598–604, Aug. 2010.

24. K. P. Weston, X. Gao, J. Zhao, K.-S. Kim, S. E. Maloney, J. Gotoff, S. Parikh, Y.-C. Leu, K.-P. Wu, M. Shinawi, J. P. Steimel, J. S. Harrison, and J. J. Yi. Identification of disease-linked hyperactivating mutations in UBE3A through large-scale functional variant analysis. Nat Commun, 12(1):6809, Nov. 2021. Number: 1 Publisher: Nature Publishing Group.

